# Understanding and mitigating some limitations of Illumina© MiSeq for environmental sequencing of Fungi

**DOI:** 10.1101/184960

**Authors:** Dan Thomas, Roo Vandegrift, Graham Bailes, Bitty Roy

## Abstract

ITS-amplicon metabarcode studies using the illumina MiSeq sequencing platform are the current standard tool for fungal ecology studies. Here we report on some of the particular challenges experienced while creating and using a ribosomal RNA gene (rDNA) amplicon library for an ecological study. Two significant complications were encountered. First, artificial differences in read abundances among OTUs were observed, apparently resulting from bias at two stages: PCR amplification of genomic DNA with ITS-region Illumina-sequence-adapted-primers, and during Illumina sequencing. These differential read abundances were only partially corrected by a common variance-stabilization method. Second, tag-switching (or the shifting of amplicons to incorrect sample indices) occurred at high levels in positive mock-community controls. An example of a bioinformatic method to estimate the rate of tag switching is shown, some recommendations on the use of positive controls and primer choice are given, and one approach to reducing potential false positives resulting from these technological biases is presented.

## Introduction

ITS or 16S amplicon libraries sequenced with Illumina MiSeq sequencing technology have become a standard tool for fungal and bacterial ecology studies. The power of next generation sequencing technologies like MiSeq, however, is balanced by its limits and biases, fueling a lively discussion on their proper implementation in microbial ecology (Pinto 2012, Lindahl 2013, Persoh 2013, McMurdie 2014 Tedersoo 2015, Nugyen 2015 and 2016, Taylor 2016). Our study continues the discussion of some of the issues surrounding metabarcoding methods, with a focus on (1) the difficulties of ecological interpretation of read abundances from Illumina-sequencer results and (2) misassignment of sample indices to reads, called *tag-switching* (alternately “index-switching” or “tag-hopping”).

Read abundances resulting from next generation sequencing studies with multiple samples and multiple biological units of interest (e.g., “OTUs”, “ESVs”. etc., etc.) are an example of a multinomial, “roll-of-the-dice” sampling process at both levels (Anders 2010, McMurdie 2014). Differences in read abundances among samples or among OTUs within samples may represent real biological differences, but they must first be tested and adjusted for the natural differences that occur when “dice are rolled.” Here we observe that initial PCR amplification of the ITS region of environmental samples of genomic DNA and Illumina-platform sequencing of the resulting libraries may both introduce this family of errors into distributions of read abundances. The variability of read abundances from next generation studies are probably most effectively modeled with negative binomial distributions (Anders 2010). Failure to adequately correct for these sources of variation could result in read distributions that give the impression of ecological patterns, such as species abundance distributions as predicted by neutral models (Baldridge 2016).

Another source of bias in next-generation sequencing studies is the erroneous assignment of sample identity to a read, or *tag-switching*. The mechanisms for this error are, to date, poorly explained and seem to vary with platform (Sinha 2017, Carlson 2012). Prescriptions for mitigating the effects of misassignment are various (Nugyen 2015 and 2016, Carlson 2012, Kong 2017).

Here we report on some of the particular challenges that result from these two sources of error, and their interaction. These were experienced while creating and using an ITS amplicon library for a fungal ecology study. Synthetic mock communities are recommended as an alternative to standard mock communities (Jusino 2016). For studies using standard mock communities, a simple method is given: observe the abundance of OTUs from mock communities in negative controls to estimate the potential frequency of tag-switching. Minimum read-abundances for observations of OTUs can then be chosen as a balance of removing as many tag-switching events as possible, while retaining as much ecological signal as possible. Additional discussion is given to some hazards and limitations of illumina MiSeq sequence data.

## Methods

The following protocols were part of an ecological study () examining landscape level patterns of leaf and wood endophytes. Leaf and wood libraries were prepared separately, and the data presented here is from the wood endophyte library This library included positive and negative controls and 91 ecological samples.

### Wood endophyte sample preparation

Wood was debarked and phloem and sapwood was collected using tools that were ethanol- and flame-sterilized between cuts. Approximately 0.5 grams of wood tissue was disrupted via bead beating using three 5 mm stainless steel beads for 3 × 30 second agitation cycles (3450 oscillations/minute), followed by an additional 30s cycle with two additional 3 mm stainless steel beads. DNA was extracted from homogenized debarked woody tissues using a Qiagen DNeasy 96 Plant Kit following the manufacturer’s instructions.

Samples were tested for presence of endophytic fungi using a preliminary PCR amplification and gel visualization of full ITS region with fungal specific primers (Gardes 1993). Successfully tested samples (91) and 3 controls were then amplified in triplicate PCRs using ITS1F forward and ITS2 reverse primers, covering the ITS1 region (Blaalid 2013), with Illumina adapter sequences and dual-indexed barcodes appended (Integrated DNA Technologies, Coralville, IA), as described above. Samples were identified using 94 unique combinations of twelve forward and eight reverse 8 bp barcodes (full primer sequences are available in the Supplemental Materials). Triplicate PCRs were performed in 20 μL volumes using the following PCR recipe: 0.6 μL each forward and reverse primers (10 μM), 0.8 μL additional MgCl_2_ (25 nM), 2.5 μL template DNA, 5.5 μL water, and 10 μL 2X PCR Super Master Mix, which contains Taq polymerase, dNTPs, and MgCl_2_ (Biotool©, now Bimake©, Houston, TX). PCR protocols: initial denaturation of 94 °C for 5 min, followed by 30 cycles of 94 °C for 30 sec, 55 °C for 1 min, and 72 °C for 30 sec each, and a final elongation of 72 °C for 7 min. Triplicate PCR products were combined and cleaned with Mag-Bind© Rxn PurePlus (OMEGA bio-tek©, Norcross, GA) beads, in equal volumes to the PCR product. Preparation of PCR plates were undertaken in a Purifier Logic+ Class II biological safety cabinet (Labconco©, Kansas City, MO).

Illumina MiSeq library preparation, after cleaning, was done using the services of the Genomics and Cell Characterization Core Facility of the Institute of Molecular Biology of the University of Oregon (Eugene, OR). Samples were normalized and pooled, along with samples from another study for a shared Illumina run. The amount of DNA being pulled from each sample was 10.45 ng (maximum allowed by the lowest concentration sample), with 258 × 10.45 ng = 2696.1 ng total, in a final volume of 384.47 μL = 7.013 ng/μL final pool concentration. Size selection was done using a Blue Pippen system with a 1.5% agarose cassette (Sage Science, Inc., Beverly, MA) to exclude DNA fragments with less than 250 bp lengths. Average ITS1 fragment length was 343 bp. Fragments larger than expected ITS1 lengths were removed bioinformatically after sequencing. Final DNA concentration within 250-1200 bp range was 5.213 nM, eluted in approximately 30 μl.

Illumina MiSeq platform sequencing occurred at the Center for Genome Research and Biocomputing at Oregon State University (Corvallis, OR) using a 600 cycle (2 × 300 bp) v3 MiSeq reagent kit and including a 10% PhiX spike-in. Quantification of the shared library using qPCR was also done at the Center for Genome Research and Biocomputing facility. Reads from the shared run totaled to approximately 23 × 10^6^ sequences, of which approximately 5.5 × 10^6^ were from the present study.

### Mock community construction

In addition to ecological samples, a pure-water negative control and two positive controls (in the form of “mock communities”, as suggested by Nguyen 2015) were included with the wood fungal endophyte library. To construct the positive controls, purified genomic DNA from 23 species of fungi from three phyla (19 Ascomycota, 3 Basidiomycota, and 1 Mucoromycota) were quantified using a NanoDrop 1000 UV-Vis Spectrophotometer (Thermo Scientific, NanoDrop products, Wilmington, DE) and diluted to a mean concentration of 9.44 ng/μl (SD = 2.35), then combined into a single sample for inclusion in the multiplexed wood fungal endophyte library. An ITS-region-only positive control was also generated using these same 23 species of fungi, using ITS1F and ITS4 primers (Gardes 1993) to amplify the full ITS region of each fungal species. PCR reactions were performed in 20 μL volumes using the following recipe: 0.6 μL each forward and reverse primers (10 μM), 0.8 μL additional MgCl_2_ (25 nM), 4.0 μL template DNA, 4.0 μL water, and 10 μL 2X PCR Super Master Mix, which contains Taq polymerase, dNTPs, and MgCl_2_ (Biotool©, now Bimake©, Houston, TX). PCR protocols: initial denaturation at 95 °C for 5 min, followed by 34 cycles of 95 °C for 1 min, 55°C for 1 min, and 72 °C for 1 min each; and a final elongation of 72 °C for 10 min. PCR products were purified with Zymo© Clean and Concentrator column kits (Zymo Research Corp., Irvine CA). Full ITS PCR product from each fungal species was then diluted to a mean concentration of 24.30 ng/μL (SD=1.74) and combined to provide a second, ITS-region-only positive control. Full ITS region PCR product from each member of the mock community were sequenced using Sanger sequencing at Functional Biosciences, Inc (Madison, Wisconsin) on ABI 3730xl instruments using Big Dye V3.1 (ThermoFisher Scientific, Waltham, MA), to provide sequence information for UNITE database taxonomy assignments (Table 1) and to provide reference sequences for downstream recovery of these fungal sequences when examining positive controls (see below). All mock communities were prepared in a physically separate location from PCR preps of ecological samples to avoid cross-contamination.

**Table 1.**
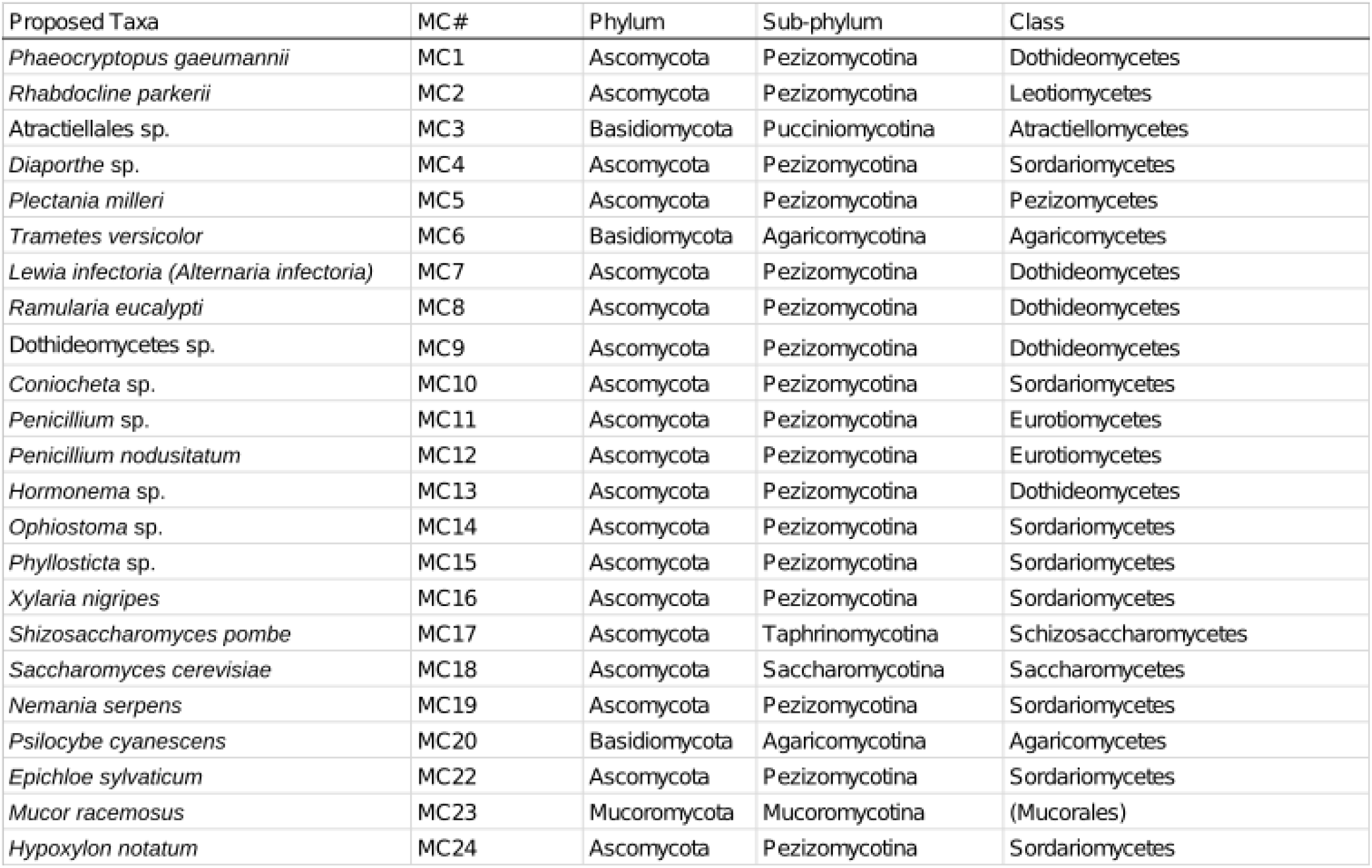
Taxa used in mock community (MC) positive control.

### Bioinformatics

General bioinformatics protocols followed the USEARCH/UPARSE pipeline version 8.1 (Edgar 2013) wherever possible. Full scripts available in supplementary information (available here and here).

FASTX toolkit software was used to visualize quality and trim unpaired read ends. After removing low quality end base-calls from each direction, paired ends were merged using the USEARCH algorithm (“fastq_mergepairs” command). Quality filtering of merged reads was implemented using the USEARCH algorithm (“fastq_filter” command) with an Expected Error approach. Primer sequences were removed from all sequences. Small numbers of reads containing “floating” primer sequences, forward and reverse primer sequences in central regions (Balint 2014), were presumed erroneous and removed using custom scripts (available in supplements). First chimera checks were conducted using the UCHIME algorithm (“uchime_ref” command) using the UNITE vers. 7.0 ITS1 reference database formatted for UCHIME. Leaf and wood libraries were concatenated at this point, and all reads were trimmed to ITS1 region only, using locations verified by the ITSx software (Bengtsson-Palme 2013). A 95% similarity radius according to UCLUST similarity algorithms in the ITS1 region was used to define OTUs. This radius was shown in our positive controls to cause less artificial splitting of fungal species (Fig. 6), while not causing noticeable artificial lumping of positive control species within the same same genus. Assignment of taxonomy to OTUs was accomplished using a modified version of the UNITE vers. 7.0 database (Kõlialg 2013): all accessions in this database not identified to at least class-level were removed. This was done to avoid the possibility that other highly probable matches with more complete taxonomic information would be ignored during taxonomic assignments. BIOM-format tables were constructed with USEARCH algorithms of the *usearch_global* program, which also allowed for inclusion of taxonomy information. Site metadata was added using the *biom-format* package (McDonald 2012). Some reformatting of taxonomic metadata of USEARCH-generated BIOM tables was required for parsing in downstream analyses. Variance stabilization of read-abundances was conducted using the *DESeq2* package in R (Love 2014. McMurdie 2013) after removal of controls.

Fungal species intentionally placed into positive control samples were distinguished from contaminants by querying with BLAST algorithm (Altschul 1990) the sequences found in our Illumina library control samples against a database of Sanger-generated sequences of the 23 intended members of our mock-community. High confidence matches were assumed to be the original, intentional members of the mock community, and the remaining sequences to be contaminants. Similarly, patterns of tag-switching were examined by querying all sequences from negative controls against this mock-community database. As most of the mock-community species were not common lab contaminants, and as care was taken during preparation of the mock community to avoid cross-contamination of Illumina libraries with DNA from positive controls before amplification of all samples with Illumina-tagged primers, the presence of members of mock-community species was interpreted as tag-switching of reads from positive control to negative control indices. The use of synthetic sequence positive control mock communities will allow for absolute, unambiguous interpretation of this pattern (Jusino 2016), and is recommended by the authors.

Artificial OTU splitting of mock-community fungal species was observed even at a 95% similarity radius for OTU formation. Due to the possible biases of PCR, OTU splitting, and tag-switching (Fig. 3), high minimum cutoffs were applied to all observations used in downstream analyses. We subtracted 60 reads, or 1.0 × 10^−5^ of total wood endophyte library size, from all observations of OTUs in each sample, and following this, observations with less than 1 read were adjusted to zero. As potential results of contamination from tag-switching, all observations of positive control fungi were removed from non-control (“ecological”) samples. After minimum abundance cutoffs and removal of any observations of mock community members in the study, 15.5% of total reads were lost. Due to this, 80.4% of observations of presence of an OTU in a sample were lost, effectively removing any information on fungi observed in low abundances.

## Results

### Positive controls

Positive controls recovered 22 of 23 species included in our mock-community. One species included in our mock communities, *Schizosaccharomyces pombe*, was not detected. A rank abundance plot of positive-control read abundances displayed a negative binomial (geometric) distribution, typical of amplicon libraries (McMurdie 2014). ITS-only positive controls displayed less dramatic differences in read abundances amoung OTUs, though large differences were still observed (Fig. 1, Fig. 2).

**Figure 1.**
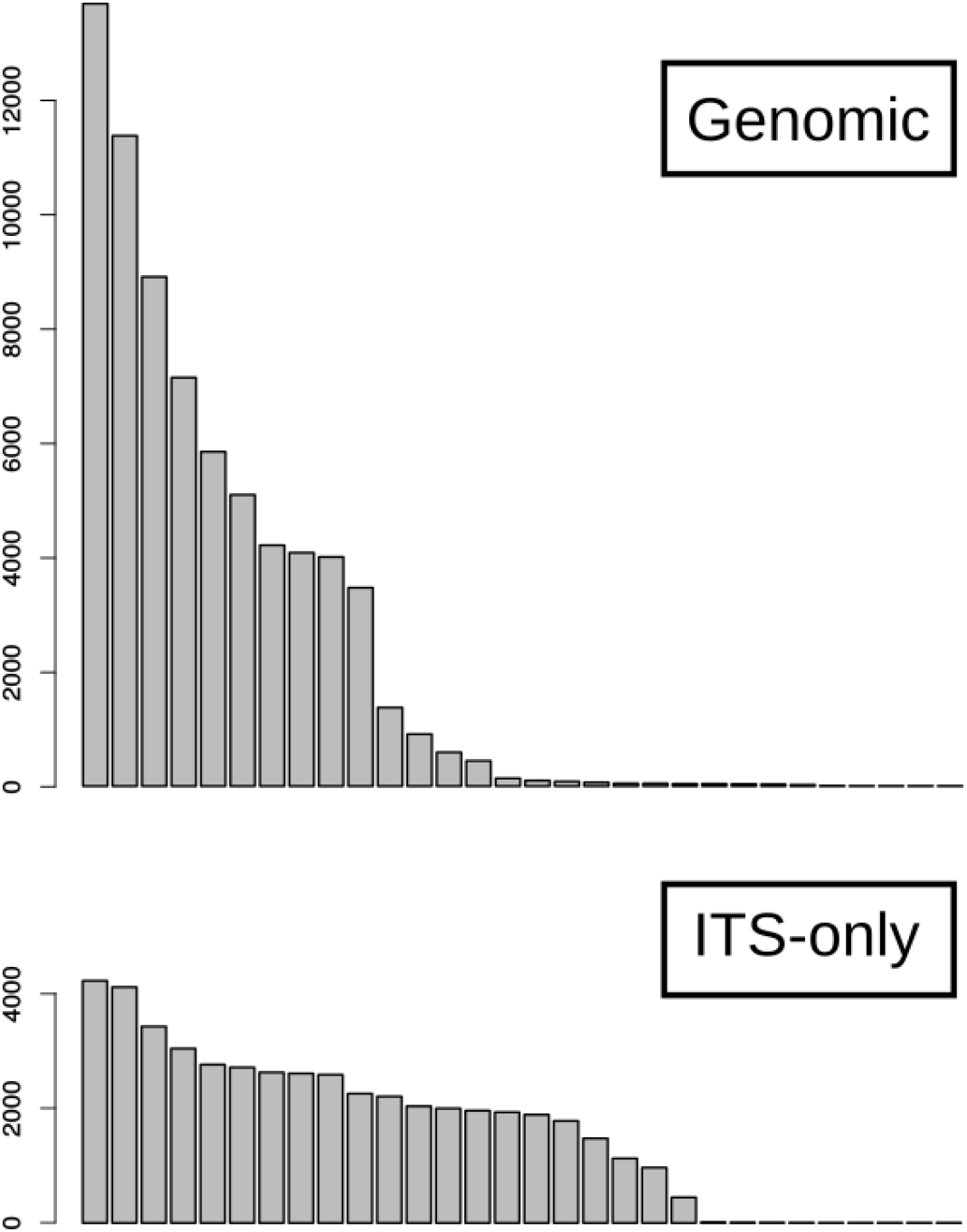
Ranked read distribution of genomic and ITS-only positive controls, by OTU. Singletons are removed after 30 OTUs. Click here for a higher resolution image.

**Figure 2.**
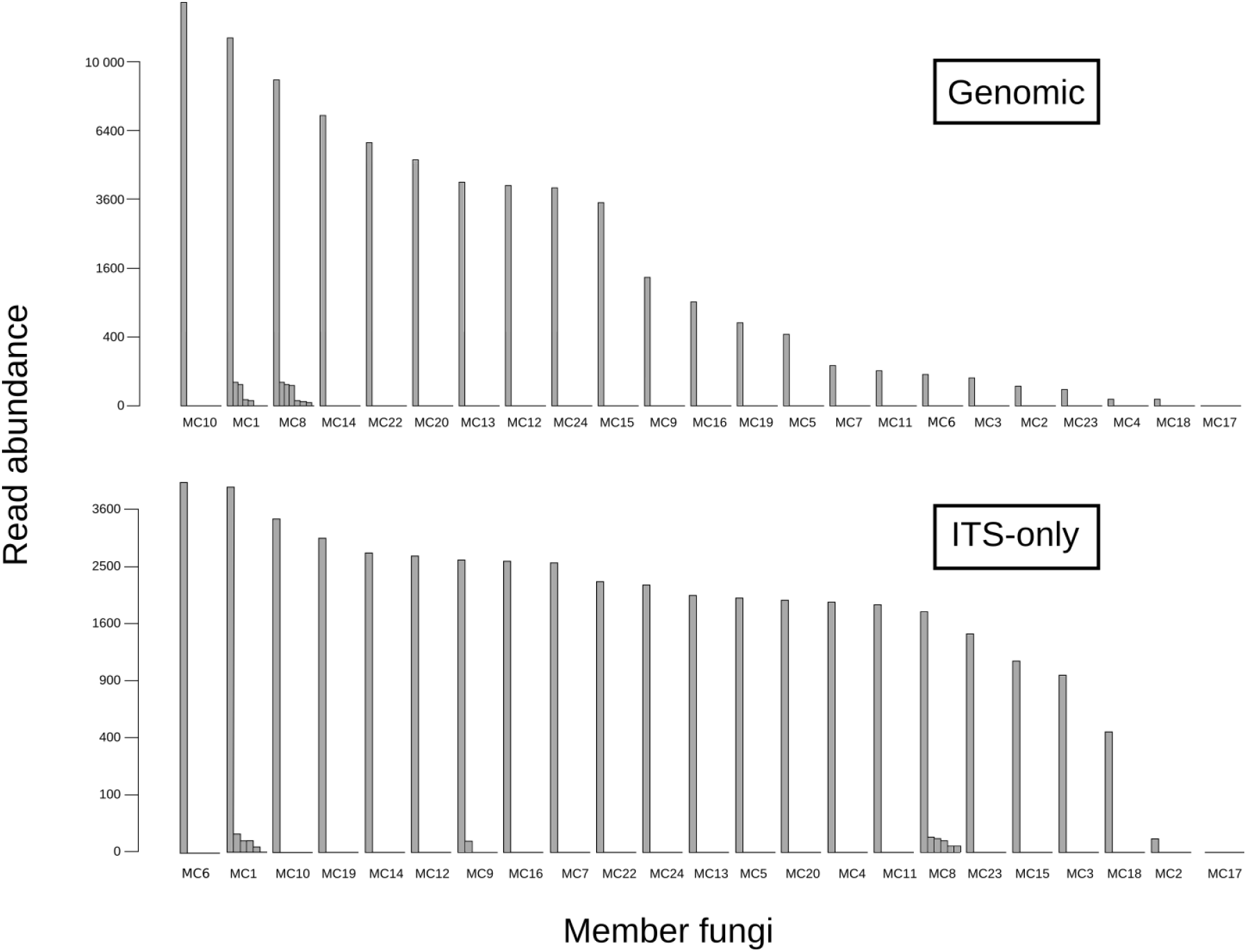
Ranked read distribution of abundances of genomic and ITS-only positive controls, including OTU splitting of mock-community members. Vertical axis is square-root transformed. Contaminants have been removed. Click here for a higher resolution image.

**Figure 3.**
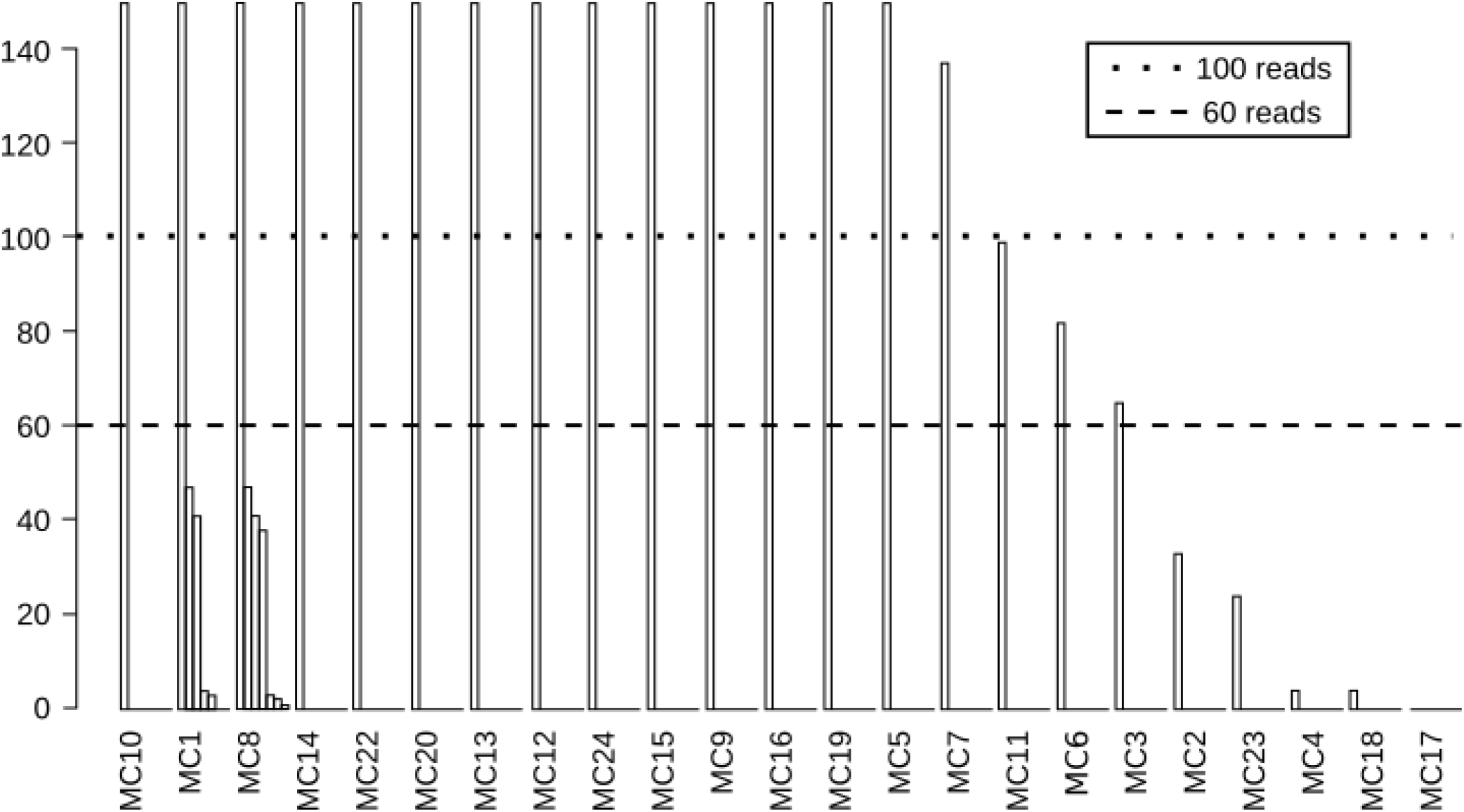
Truncated ranked read distribution of abundances of genomic positive control, including OTU splitting of mock-community members. The 100-read line represents the level around which tag-switching errors were observed to occur in the study (see fig. 7), the 60-read line represents the abundance that authors chose as a minimum abundance cut-off for observations. Click here for a higher resolution image.

### Variance stabilization

Transformation by *DESeq2* algorithms adjusted total read levels to more equal proportions among all samples (fig. 4), and reduced the scale of artificial differences from PCR/sequencing bias among read abundances of OTUs within our positive controls (fig. 5), and therefore also presumably in ecological samples (fig. 6). Despite this, read differences of one order of magnitude were found among our genomic mock-community samples even after variance stabilization (Fig. 5).

**Figure 4.**
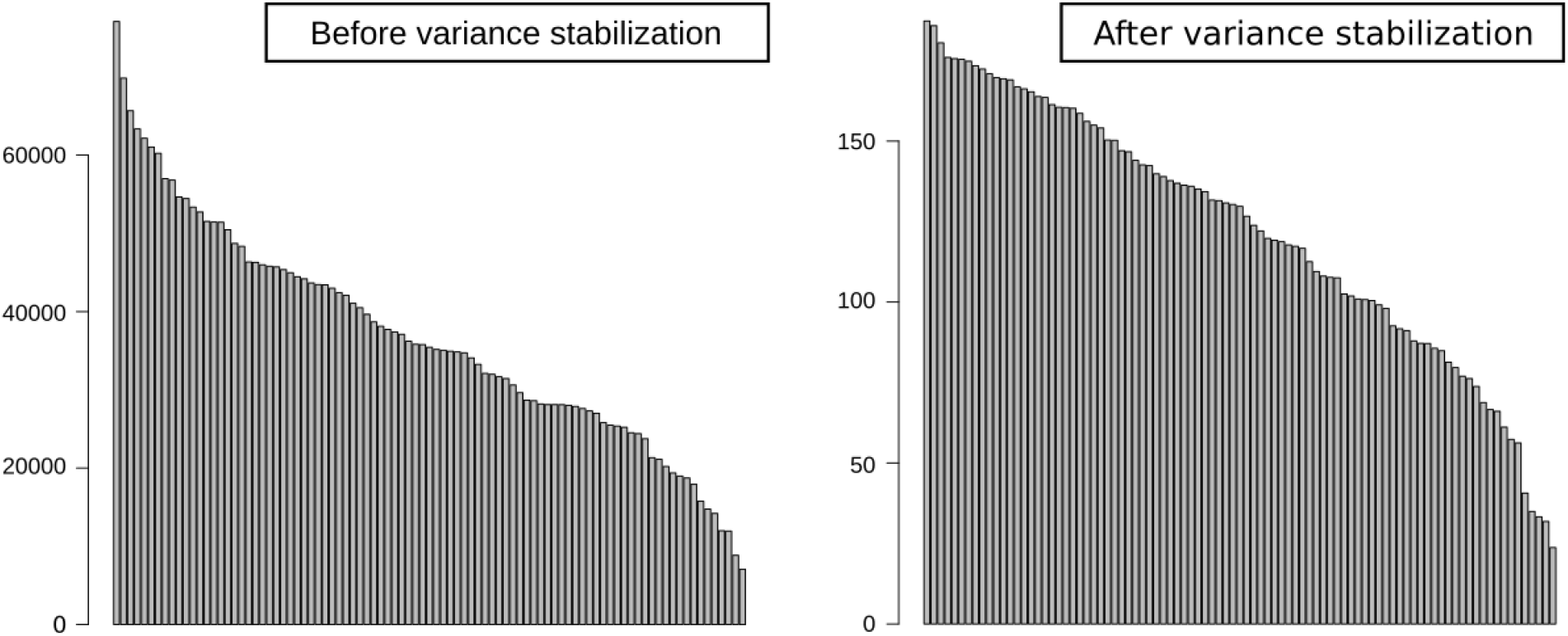
Ranked distribution of read abundances per sample for entire wood endophyte library, before and after variance stabilization using DEseq2 algorithms. Click here for a higher resolution image.

**Figure 5.**
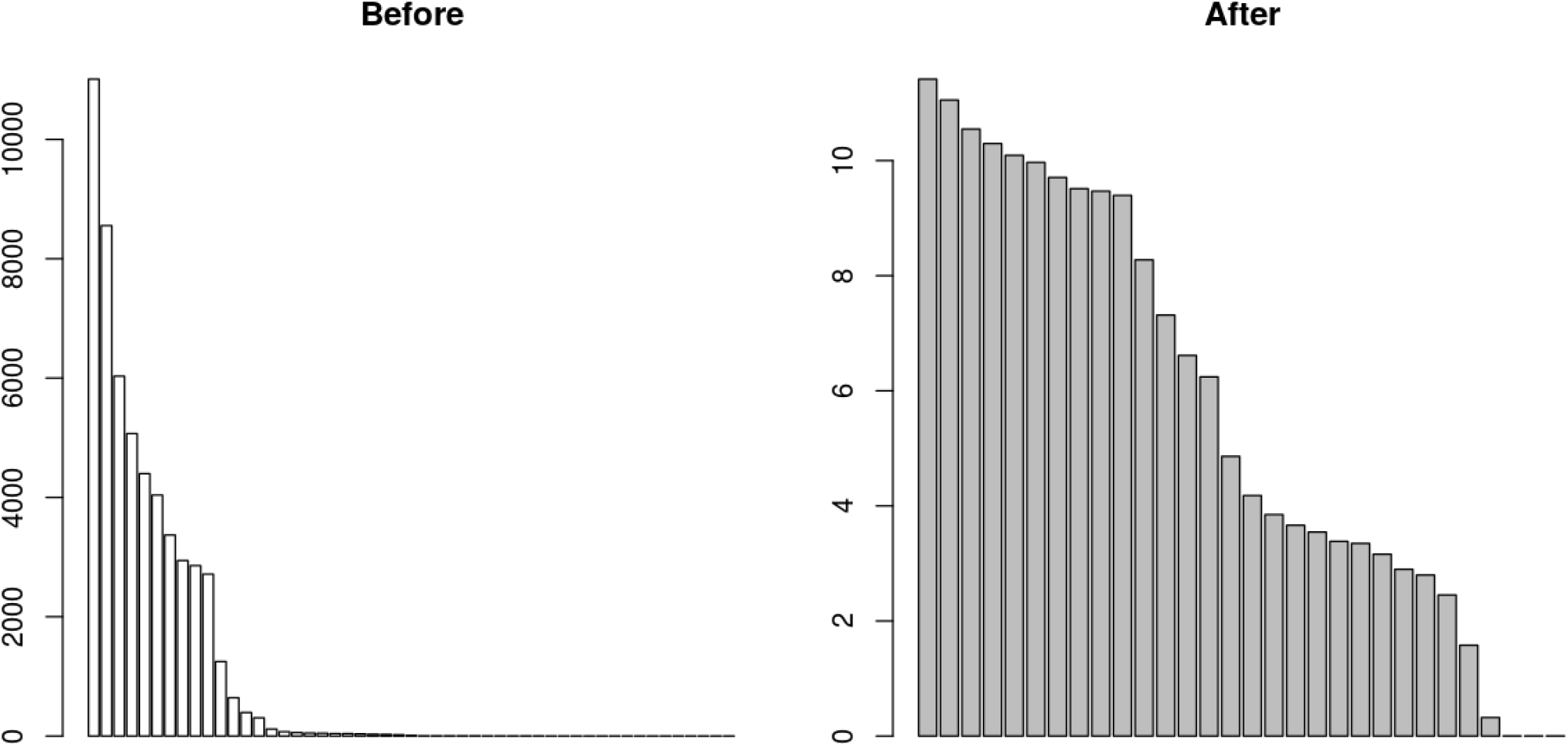
Ranked distribution of read abundances per OTU for the genomic positive control, before and after variance stabilization using *DEseq2* algorithms. Click here for a higher resolution image.

**Figure 6.**
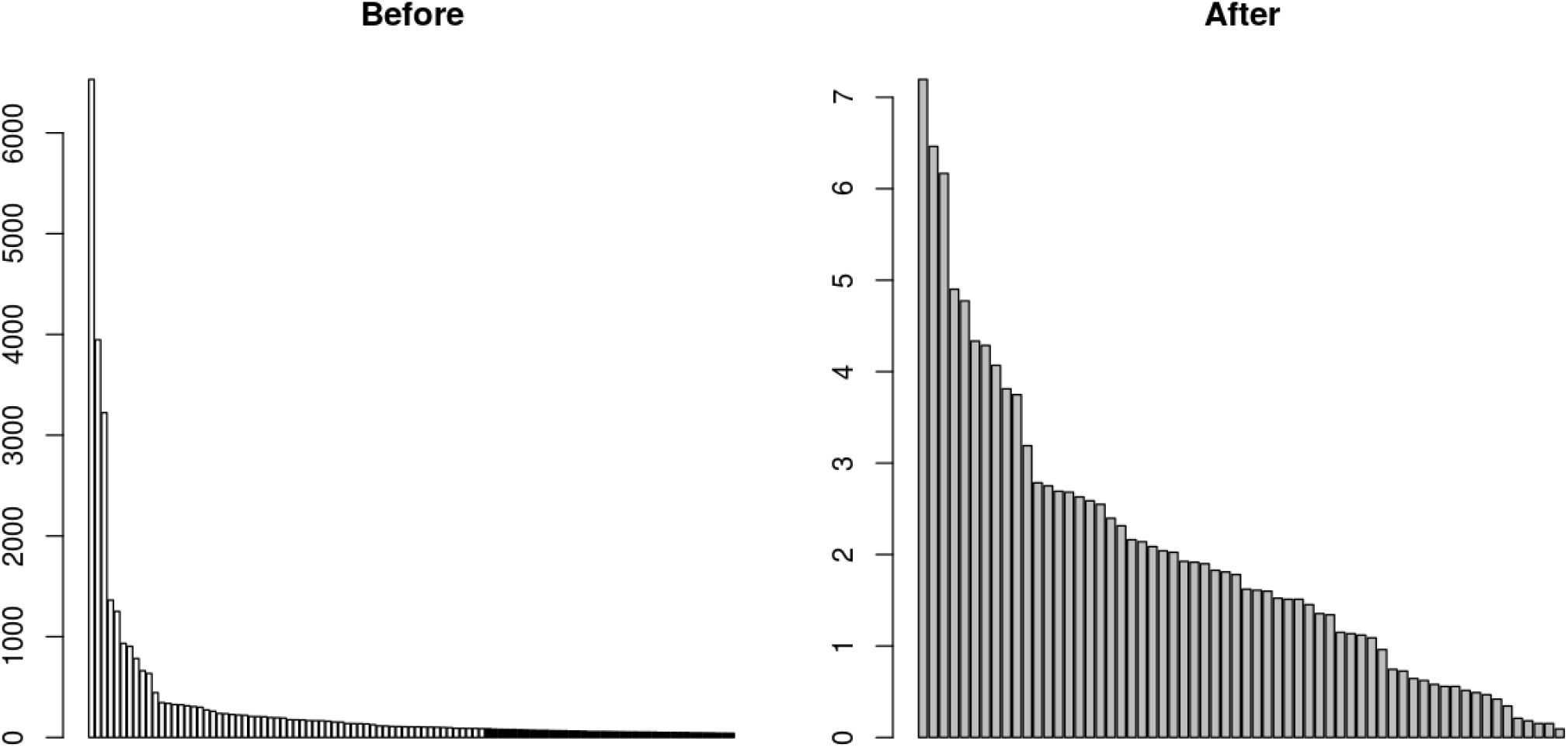
Ranked distribution of read abundances per OTU for one randomly selected sample, before and after variance stabilization using *DEseq2* algorithms. Blackened OTUs (left) represent OTUs that are removed by variance stabilization. Click here for a higher resolution image.

### Negative controls

Using an OTU similarity radius of 95%, pure water control contained 54 OTUs, with abundances of individual OTU observations up to 544 reads. 13 of these OTUs present in negative controls matched with high confidence to members of our positive control communities (Fig. 7).

**Figure 7.**
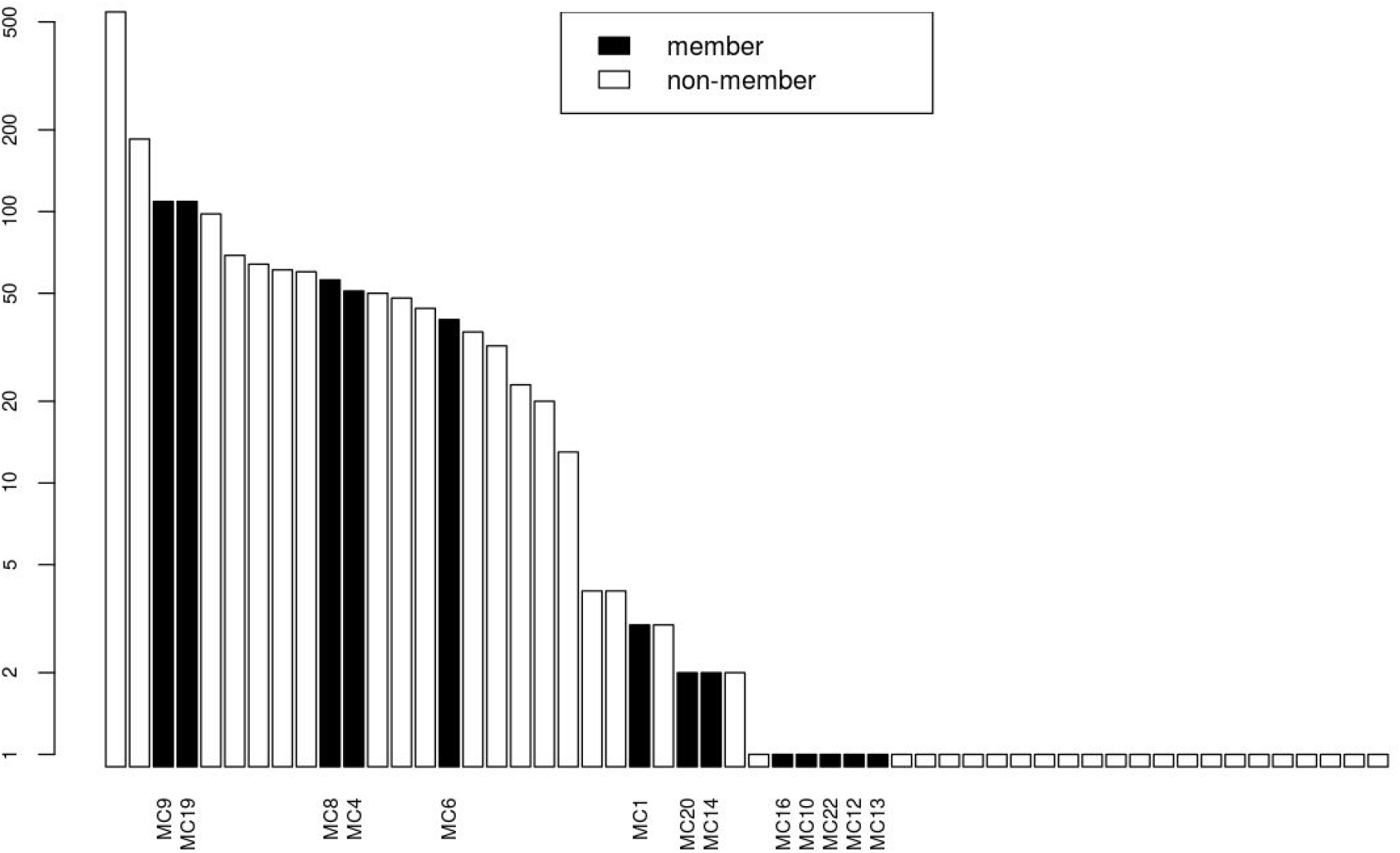
Ranked read distribution of OTUs from a pure water negative control. Black bars indicate OTUs that are also members of positive control, indicating probable misassignment of reads. Click here for a higher resolution image.

## Discussion

The approximately negative binomial curve of genomic controls, compared to the less dramatic differences in abundances observed in our ITS-only positive controls (Fig. 5), suggests that much of the potential bias within this study, and potentially amplicon sequencing studies in general, may originate in the initial PCR step. ITS regions of organisms must be “found” amid the other regions of many genomes of the thousands of organisms present in any environmental sample of DNA. Initial conditions that allow easier discovery, such as larger ITS copy numbers (Schoch 2012) or ease of DNA extraction (Fredericks 2005), may be very important in determining which organisms’ barcode regions are ultimately represented in amplicon sequence libraries. Even after adjusting for these differences among samples through negative-binomial variance stabilization, large artificial differences in read abundances remained within our genomic positive controls.

Our pure-water negative control showed high levels of sequences likely to have originated from our positive control mock-community. These patterns of tag-switching indicate that mock-community DNA disproportionately affected other samples in the study, probably due to relatively high concentrations of the simpler mock community DNA as compared to diverse ecological samples.

To utilize the data presented here in downstream ecological analyses (Thomas 2034), we chose to use a presence/absence transformation of data to correct for artificial differences in read abundance. This method results in the elevation of low-abundance observations of OTUs to equal importance with higher abundance observations in downstream statistical analyses. This was deemed appropriate given that even after variance stabilization, large artificial differences in abundances remained in the positive control. This elevation of importance for low-abundance OTUs can be problematic, as low-abundance observations and rarely observed OTUs are more likely to be spurious (Huse 2010, Brown 2015): here OTU splitting and tag-switching caused low-abundance, erroneous OTUs that were present as more than singletons (Fig. 3). Mock-community samples were used to estimate levels of tag-switching and generate minimum read-abundances for observations, as well as gauge appropriate levels of similarity for definition of OTUs, and to gain insight into levels of PCR amplification biases. If tag-switching is shown to be common in an amplicon study, and presence/absence transformation is used to correct for artificial differences in read abundances, higher minimum cutoffs per observation than traditional removal of singleton OTUs are appropriate to avoid the elevation of spurious OTUs to the same importance as real OTUs. This, of course, results in a large loss of information about the presence of rare organisms (Brown 2015) (Fig. 6); in this study, strong ecological signals remained even after presence/absence transformation and application of strict minimum read abundance cutoffs (Thomas 2034).

Illumina has stated that index misassignment occurs at low levels and is likely due to contamination from free, unligated adapter/primer oligonucleotides (Illumina 2017). In the present study however, free primers should have been largely removed at two stages in library preparation: cleaning of all PCR products with Mag-Bind beads and size selection of fragments larger than 250 base-pairs. Despite this, erroneous, tag-switched reads appear at levels equal to the rate at which of many “real” OTUs appeared in the positive control (Fig. 3, Fig. 7). Thus, dismissing the issue of tag-switching due to the relatively low rate of index misassignment misses the mark for microbial metabarcoding studies: the researcher is confronted with balancing the loss of real ecological information against the need to remove possible incidences of tag-switching that could create false positives. In this study, after bioinformatic processing, 7 of the 22 species present as OTUs in the mock community were present below the 100 base-pair abundance. This 100 base-pair abundance is the level at which some putatively tag-switched OTUs were observed in the negative control (Fig. 3, Fig. 7).

The confusion of OTU observations with errors resulting from tag-switching is compounded if species of interest are included in the positive control. Contrary to the recommendations of Kong et al. (2017), researchers should also take care to avoid the use of species of biological interest as members of their mock communities, as observations of these species outside of positive control samples may then be called into question as a potential result of tag-switching, particularly where positive control mock communities have high overall DNA concentrations. In the present study, significant levels of tag-switching were evident, especially for the members of the positive control mock community itself (Fig. 7), so all OTUs identified as original members in the positive controls were removed entirely from the library for downstream analyses. This study highlights the need for strict cutoffs and careful implementation of positive controls, and a framework for estimating rates of tag-switching from these. The most promising toolset for estimating rates of tag-switching is a completely synthetic positive control proposed by Jusino et al. (2016), which is a mock-community constructed from fungus-like ITS-region oligonucleotides that do not represent any organisms in nature.

